# Formal meta-analysis of hypoxic gene expression profiles reveals a universal gene signature and cell type-specific effects

**DOI:** 10.1101/2021.11.12.468418

**Authors:** Laura Puente-Santamaria, Lucia Sanchez-Gonzalez, Barbara P. Gonzalez-Serrano, Nuria Pescador, Oscar H. Martinez-Costa, Ricardo Ramos-Ruiz, Luis del Peso

## Abstract

**Background:** Integrating transcriptional profiles results in the identification of gene expression signatures that are more robust than those obtained for individual datasets. However, direct comparison of datasets derived from heterogeneous experimental conditions is not possible and their integration requires the application of specific meta-analysis techniques. The transcriptional response to hypoxia has been the focus of intense research due to its central role in tissue homeostasis and in prevalent diseases. Accordingly, a large number of studies have determined the gene expression profile of hypoxic cells. Yet, in spite of this wealth of information, little effort have been done to integrate these dataset to produce a robust hypoxic signature.

**Results:** We applied a formal meta-analysis procedure to a dataset comprising 425 RNAseq samples derived from 42 individual studies including 33 different cell types, to derive a pooled estimate of the effect of hypoxia on gene expression. This approach revealed that a large proportion of the transcriptome (8556 genes out of 20888) is significantly regulated by hypoxia. However, only a small fraction of the differentially expressed genes (1265 genes, 15%) show an effect size that, according to comparisons to gene pathways known to be regulated by hypoxia, is likely to be biologically relevant. By focusing on genes ubiquitously expressed we identified a signature of 291 genes robustly and consistently regulated by hypoxia. Finally, by a applying a moderator analysis we found that endothelial cells show a characteristic gene expression pattern that is significantly different from other cell types.

**Conclusion:** By the application of a formal meta-analysis to hypoxic gene profiles, we have developed a robust gene signature that characterizes the transcriptomic response to low oxygen. In addition to identifying a universal set of hypoxia-responsive genes, we found a set of genes whose regulation is cell-type specific and suggest a unique metabolic response of endothelial cells to reduced oxygen tension.

## Introduction

Oxygen homeostasis is essential to sustain cellular metabolism in eukaryotes. Hypoxia triggers multiple adaptive mechanisms, from metabolism reprogramming to tissular restructuring, aimed to re-balancing oxygen supply and demand [1]. In multicellular organisms this response can be very diverse, depending on cell type, extension and degree of the oxygen deprivation, or pathological state.

Most of these responses are orchestrated at the transcriptional level, with the Hypoxia Inducible Factors (HIFs) being the main drivers of the hypoxic gene expression pattern [2]. The heterodimeric HIF transcription factor consists on a *β* subunit (ARNT), constitutively expressed, and an *α* subunit (HIF1A, EPAS1, HIF3A) which, in normoxic conditions, is marked for degradation by the concerted action of a family of oxygen-dependent enzymes (EGLN family) and the von Hippel-Lindau (VHL) ubiquitylation complex [3–5]. When oxygen concentration decreases, the *α* subunits escape degradation, due to the reduced activity of the EGLNs, and are translocated to the cell’s nucleus, where they bind to Hypoxia Response Elements along the *β* subunit. Transcriptional activity of HIFs depends also on interaction with co-activators such as CREB-binding protein or p300 whose binding is also regulated in an oxygen-dependent manner [6, 7].

Given the importance of the transcriptional response for tissue oxygen homeostasis and its alteration in disease, a large number of works have attempted to identify the full set of genes regulated by hypoxia by means of gene profiling experiments. Since these studies were performed in a wide variety of experimental conditions including different cell types, oxygen tensions and exposure times, the integrated analysis of their results could be exploited to identify a set of genes ubiquitously regulated by hypoxia as well as set of genes whose regulation by hypoxia is restricted to specific situations. However, little effort has been done in this regard and, to the best of our knowledge, only two attempts to integrate all the hypoxic gene profiling experiments have been done [8, 9]. The first analysis of this type, based on the analysis of gene profiles generated by means of DNA microarrays, produced the first list of genes universally induced by hypoxia and revealed that the set of genes induced by hypoxia were more conserved than those repressed by hypoxia [8]. A second, more recent study, exploited the information derived from RNAseq experiments producing a more comprehensive list of hypoxia-regulated genes and characterized HIF-isoform common and specific targets [9]. In spite of their merit, none of this works employed formal meta-analysis approach for their analysis which, given the heterogeneous nature of the data, is critical to draw statistically sound conclusions [10]. In particular, in a random-effects meta-analysis model, the effect sizes derived from individual studies are assumed to represent a random sample from a particular distribution of these effect sizes (hence the term random effects). In other words, instead of assuming a fixed effect of hypoxia on any given gene across the different studies, under the random effects model we allow that the true effect could vary from study to study to reflect for example the different response in distinct cell types. In this study we aim to define core components of the transcriptional response to hypoxia taking advantage of the wider public availability of next generation sequencing data, RNA-seq in particular. Applying a random effects model to the expression data gathered we are able to define a molecular signature representing the early (<48h) transcriptional response to hypoxia, independently of cell type. Given the abundance of studies testing the effects of hypoxia on endothelial cells we have additionally examined cell-type specific alterations of gene expression.

## Materials and Methods

### 0.1 RNA-seq data download and processing

Raw reads of the RNA-seq experiments were downloaded from Sequence Read Archive [11]. Pseudocounts for each gene were obtained with salmon [12] using RefSeq [13] mRNA sequences for human genome assembly GRCh38/hg38 as reference.

Differential expression in individual subsets was calculated with the R package DESeq2 [14] using local dispersion fit and apeglm [15] method for effect size shrinkage.

### 0.2 Meta-analysis

The meta-analysis intended to identify the effect of hypoxia on early gene expression in human cells compared to normoxic controls. To identify studies to be included in the meta-analysis Gene Expression Ommibus (GEO) repository was searched with the terms ‘hypoxia[Description] AND “expression profiling by high throughput sequencing”[DataSet Type]’ on february 11, 2021. The search resulted in a total of 394 studies. We only kept studies performed in human cells that determined steady-state RNA levels in total (poly-A) RNA samples and excluded analysis that did not include replicates, employed treatments other than reduced oxygen tension (e.g. chemical inhibitors or other hypoxia mimetics) or those where gene expression was analyzed after 48h. A total of 46 studies (independent GSE entries) remained after application of the inclusion/exclusion criteria and were used for the meta-analyses (supplementary table S2). A pooled estimate of the size effect of hypoxia on expression was determined for each gene using the R packages metafor [16] and meta [17] using as input the log2-Fold change value and its associated standard error computed for each individual RNA-seq experiment using the R package DESeq2 [14]. Given that the individual estimates derive from an heterogeneous group of experiments, including different cell types and experimental conditions, we assumed that these individual estimates derive from a distribution of true effect sizes rather than a single one and thus applied a random-effects model for the meta-analysis. As estimator of the between-study heterogeneity we used the Maximum Likelihood (“ML”) method.

Subgroup analysis (Moderator analysis) was used to compare the pooled estimated effect in studies grouped according to cell type analyzed.

### 0.3 Functional Enrichment Analysis

Enrichment of Gene Ontology terms was performed with the Bioconductor’s clusterProfiler package [18] using a q cut-off value of 0.05. To reduce the redundancy, highly similar GO terms were removed keeping a single representative by using the “simplify” function using a cut-off value of 0.6 (upregulated genes) or 0.7 (downregulted genes). The much larger number of enriched terms found for upregulated genes justified the use of a more slightly more lenient cutoff value for the simplify function.

## Results

### Meta-analysis strategy

In order to identify genes consistently regulated by hypoxia across a wide range of cell types and experimental conditions, we integrated the results from 46 studies analyzing the transcriptional response to hypoxia by means of RNA-seq (supplementary table S2). Since some studies included several cell types, oxygen tensions or times of exposure to hypoxia, we took subsets of the study’s data so that each one included a single cell line and set of experimental conditions (figure 1). Thus, our initial data set included a total of 81 subsets of normoxia-hypoxia paired samples, each one comprising a single cell line, exposure time and oxygen tension (table 1).

**Table 1.**
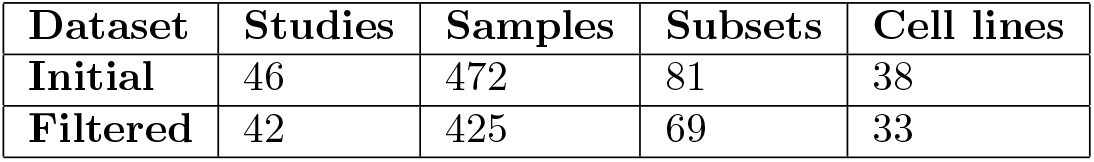
Datasets used in the meta-analysis. The number of independent GEO entries (Series IDs, GSE) that met the criteria described under methods is shown (“Studies”). For each study, only those samples corresponding to normoxic and hypoxic conditions were retrieved ignoring any other treatment that the study could have included. The total number of samples meeting these criteria is shown (“Samples”). In those cases were a study included more than one cell line, different degrees of hypoxia or different exposure times, the samples were divided into subsets including a single level for each one of these variables. The total number of subsets generated is shown (“Subsets”). The column “Cell lines” indicates the number of different cell lines included in the dataset. The rows contain values corresponding to the dataset prior (“Initial”) and after (“Filtered”) filtering to remove outlier studies.

**Figure 1.**
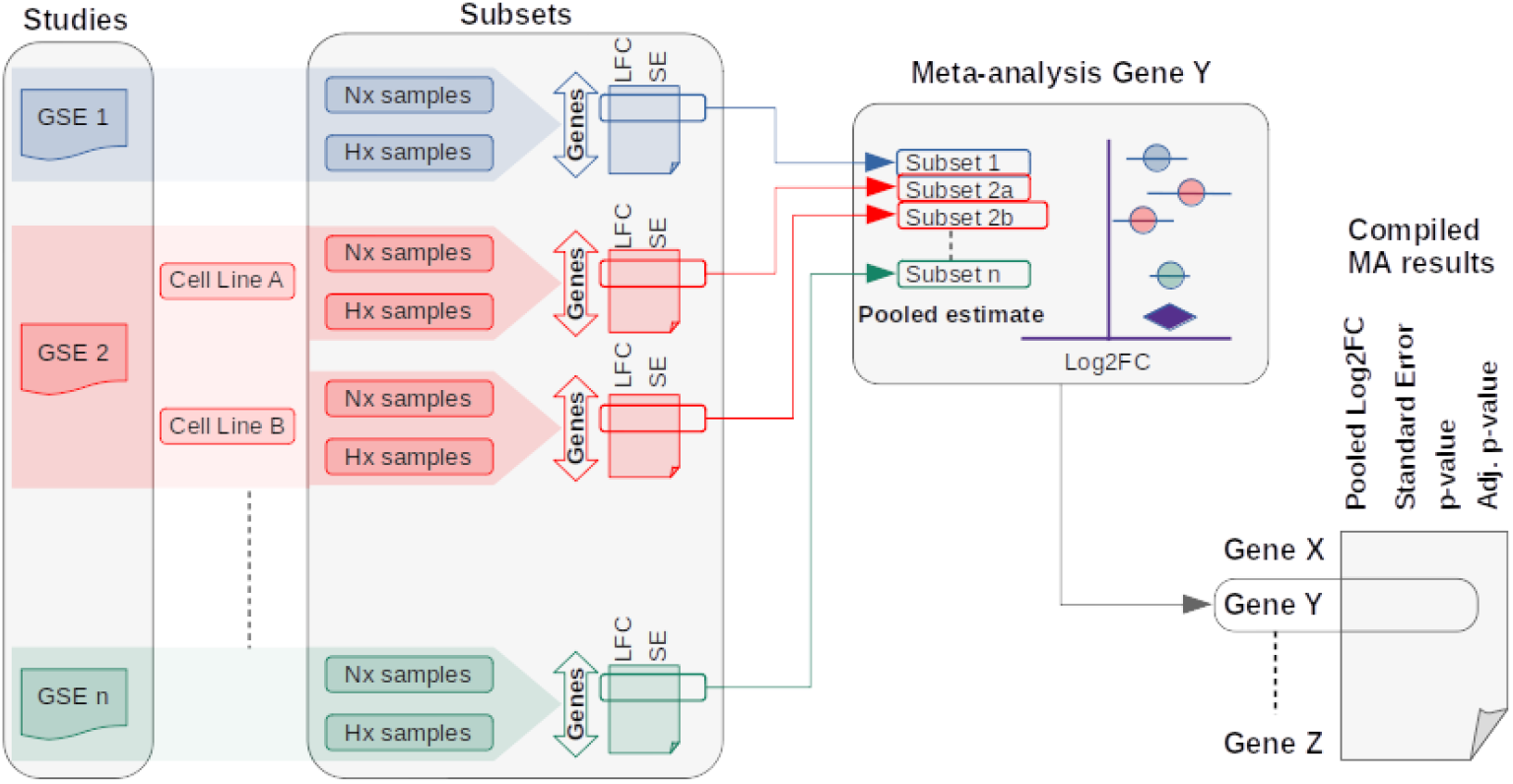
Integration of studies and gene-level meta-analysis. Normoxic (Nx samples) and hypoxic (Hx samples) replicates from the relevant studies (“GSE1”, “GSE2”,…“GSEn”) were processed to produce a table recording the effect of hypoxia on the expression of each gene (Log2 fold-change,labeled as “LFC”) and the standard error associated to this estimation (“SE”). In those studies analyzing more than a single cell line, time of exposure to hypoxia or oxygen tension, samples were grouped to generate homogeneous subsets and the effect of hypoxia on gene expression was analyzed in each individual subset. The figure represents this situation in the case of GSE2 (shaded in red color), an hypothetical study that analyzed the effect of hypoxia in two different cell types. Then, the results obtained for each individual gene were integrated into a random-effects model meta-analysis to produce a pooled estimate of the effect of hypoxia on each gene. Note that a meta-analysis is performed for each individual gene. Finally, the results of the gene-level meta-analyses were integrated into a single list (“Compiled MA results”).

Next, for each of these 81 subsets, we estimated the difference in the level expression between normoxia and hypoxia conditions for all genes (“LFC”, figure 1). From these analyses we extracted the information corresponding to individual genes (figure 1; Log2 Fold-change, “LFC”, and the standard error associated to this estimate, “SE”) across the different studies and performed a formal meta-analysis on each one of these gene-specific datasets to estimate the pooled effect of hypoxia across studies (figure 1 Meta-analysis). Thus, an independent meta-analysis was performed for each individual gene by integrating the effects of hypoxia on that particular gene across the different studies and conditions. The results provide a pooled estimate of the effect of hypoxia on the expression of the gene under analysis and its statistical significance. As an example, the results of the meta-analysis for the EGLN3 gene, encoding a cellular oxygen sensor known to be directly regulated by HIF in response to hypoxia ([19]), are shown in supplementary figure S1. Finally, we compiled the pooled estimates for all genes detected in more than one subset, together with the statistical significance value, to produce a table representing the overall effect of hypoxia on gene expression (figure 1 “Compiled MA results”).

In order to refine the results of the meta-analyses, we sought to identify outlier studies. To this end, we calculated the correlation of the effect of hypoxia across all genes in each individual study compared to the estimated pooled effect derived from the meta-analyses. As shown in figure 2A, this analysis revealed a few incoherent sets. These same subsets were identified as outliers due to their low or negative correlation with the rest of subsets in pair-wise comparisons 2B. In those cases where the lack of positive correlation was clearly due to a mistake in the labeling of samples in GEO, as indicated by a large negative correlation coefficient (figure 2, subsets “S42” and “S58”), the treatment labels were correctly set and the study was kept. The remaining incoherent studies, having a correlation coefficient not significantly different to zero (*p* > 0.01), were discarded. After these data sanity check procedures, the whole analysis strategy (1) was repeated on this corrected and filtered dataset. Table 1 shows the statistics of the data set after filtering and supplementary table S3 the full description of each of the samples included in the final analysis.

**Figure 2.**
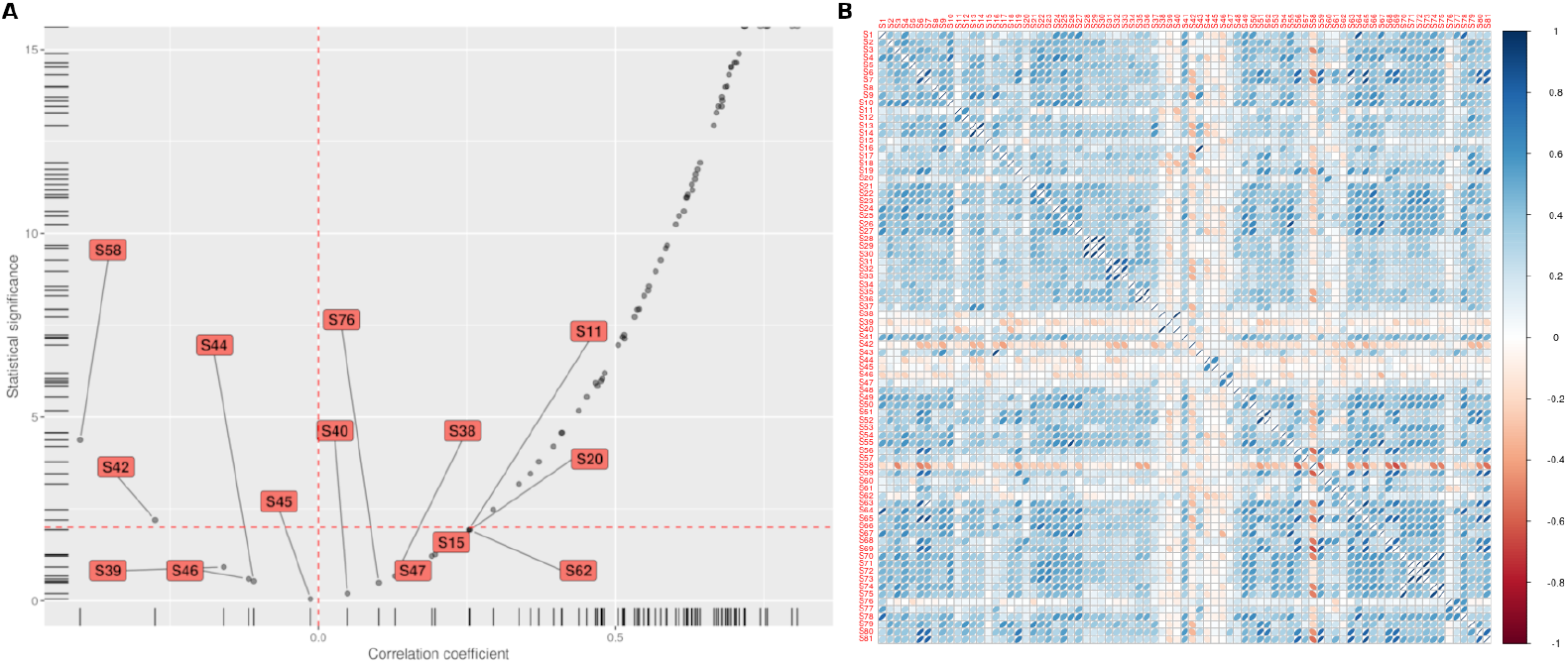
Identification of outlier subsets. A) The pooled effect of hypoxia on gene expression was compared to the estimates obtained in each individual subset for all the genes that were ubiquitously expressed (expression detected in more than 75% of subsets) and that showed a strong (absolute size effect > 1) a significant (FDR < 0.01) response to hypoxia according to the meta-analyses. Each dot represents the Pearson’s correlation coefficient (r) and its statistical significance (FDR-adjusted p-value). The horizontal and vertical dotted lines correspond to an adjusted p-value of 0.01 and a correlation coefficient of zero respectively. Those subsets that did not showed significant correlation with the meta-analyses pooled estimates (FDR > 0.01) or showed a negative correlation (r < 0) are identified and labeled. B) Correlation between all pairs of datasets. The figure shows the Pearson’s correlation coefficient for all pair-wise comparisons. The color and shape of the ellipses indicate the strength of the correlation (value of Pearson’s r) and its direction.

### Identification of a universal core of hypoxia-inducible genes

The results of the meta-analysis on the clean dataset, after filtering out the outlier subsets and removing genes detected in less than 5% of the subsets, revealed 8556 genes (out of a total of 20888) whose expression was significantly (*FDR* < 0.01) altered in response to hypoxia (figure 3A and supplementary table S4), with similar number of genes being induced (4481) and repressed (4075). These numbers are larger than the typical values obtained in individual experiments, with median values around 1000 genes in each category (Figure 3B), which is expected from the increased power to detect small size effects due to the integration of a large number of samples in the meta-analysis. In agreement, the median effect size (LFC) observed for the genes differentially expressed (DE) according to the meta-analyses are −0.29 and 0.4 for down- and up-regulated genes respectively, contrasting with the median effect size observed in individual studies of −0.72 and 0.83 for up- and down-regulated genes respectively (figure 3C).

**Figure 3.**
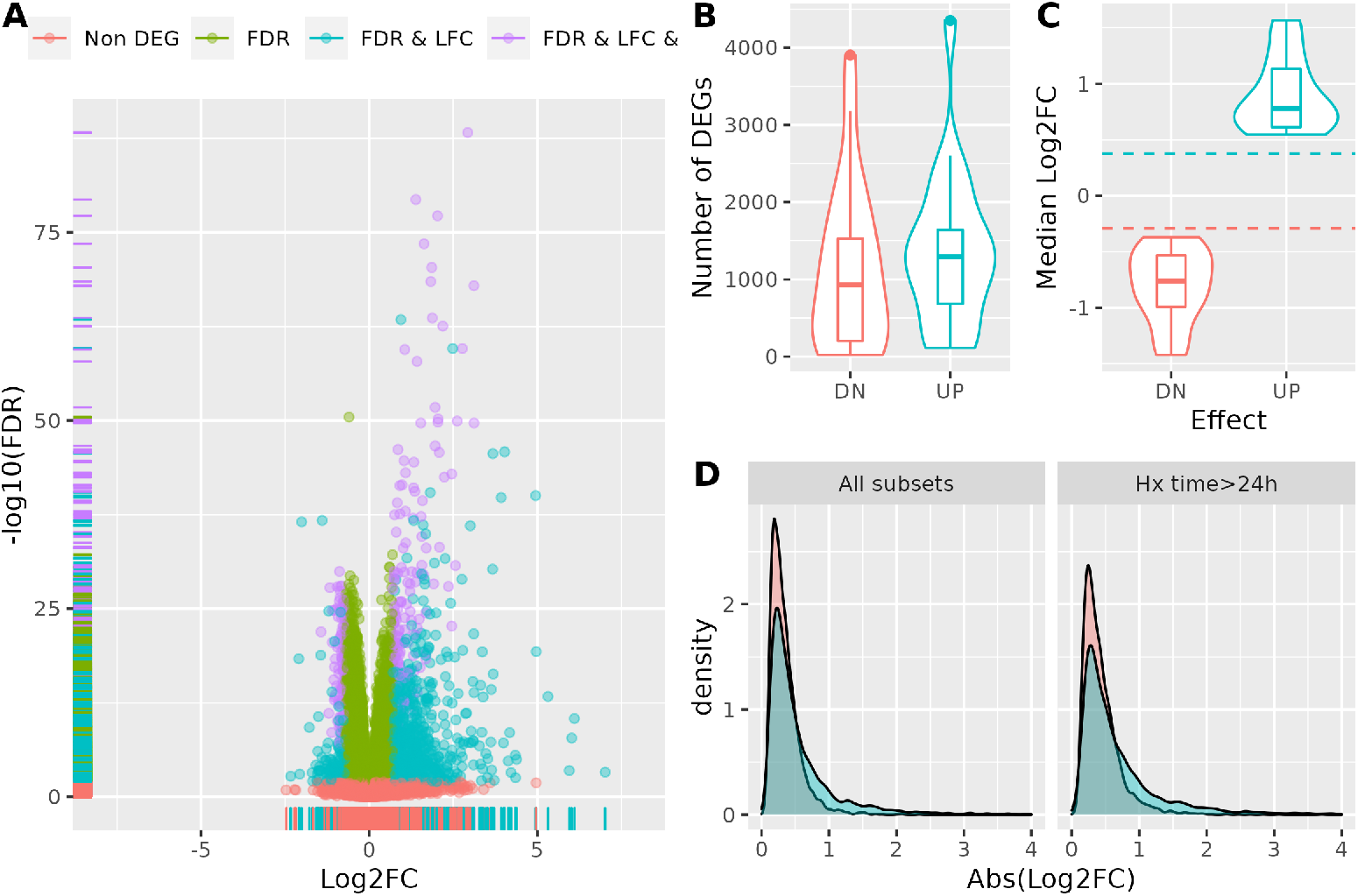
Identification of a common set of hypoxia-regulated genes.. A) The graph represents the pooled effect of hypoxia on gene expression (Log2FC hypoxia over normoxia) against the statistical significance of the effect (-log10 FDR-adjusted p-value) according to the meta-analysis. Genes are represented as dots and their color indicates the effect of hypoxia: blue, genes not regulated by hypoxia (*FDR – adjustedp – value* ≥ 0.01); green, genes mildly affected by hypoxia (*FDR – adjustedp – value* < 0.01 and │*Log*2*FC*│ < 1); red, genes robustly regulated by hypoxia (*FDR – adjustedp – value* ≥ 0.01 and │*Log*2*FC*│ ≥ 1). The y-axis was limited to a value of 65 for representation purposes (a single gene was left out). B) Distribution of the number of genes found significantly (*FDR – adjustedp – value* ≥ 0.01) down- (“DN”) or up-regulated (“UP”) by hypoxia in individual studies. c) Distribution of the median effect size (Log2FC Hypoxia over Normoxia) of hypoxia on repressed (“DN”) or induced (“UP”) genes in individual studies. The red and blue dotted lines correspond to the median effect size for repressed and induced genes respectively, according to the meta-analysis pooled estimates.

In the view of these results we tried to identify an minimum effect size likely to represent biologically relevant changes in gene expression. To this end, we compared the proportion of genes belonging to biological processes known to be regulated by hypoxia, among the genes selected according to increasing effect size (Log2 FC) cut-off values. As shown in supplementary figure S2, the p-value for the association of biological function and regulation by hypoxia reached a minimum at effect size (Log2FC) values between 0.3 and 1.7. Since the choice of ES values only affects the distribution of genes into categories (i.e.differentially expressed versus not altered) but does not change their total number, the minimum p-value corresponds the least likely distribution expected by chance. Thus, we decided to take these values as the lower boundary required to produce a biological response to hypoxia. The median value of the effect sizes is 0.7, corresponding to an induction of 1.6 times over basal levels (0.6 times normoxic level in the case of repressed genes).

Thus, in response to hypoxia a total of 1265 genes, 1020 induced and 245 repressed, show a statistically significant change in expression (*FDR* < 0.01) of a magnitude likely to be biologically meaningful (│*Log*2*FC│* > 0.7) (figure 3A labeled in purple and blue).

The difference in the number of repressed and induced genes is a consequence of the distribution of effect size values having a longer tail in the latter case (rug plot of the x-axis in figure 3A and figure 3D left panel). The different shapes of the distribution of effect size values also suggest that hypoxia has a relatively weak effect on gene repression. In fact, while the number of significantly induced genes is about five times higher that of repressed genes (1020 vs 245) for an effect size higher than 0.7, the ratio increases to seventeen times more up-regulated than down-regulated genes (268 vs 15) for effect sizes larger than 1.5. Since the meta-analyses included experiments done at relatively short exposure times (27% of the subsets correspond to exposure times ranging from 1-12h), it could be argued that the smaller effect size observed for repressed genes is a consequence of short-time experiments failing to detect the effect on mRNA levels due to their half-life. To test this hypothesis, we repeated the meta-analyses selecting only subsets corresponding to treatments of 24-48h, significantly longer than the median half-life of 5.7 h observed for human mRNAs under hypoxia [20]. As shown in figure 3D left panel, both distributions show a small shift toward higher absolute effect size values, but the difference between them remains unaltered. Thus, the relatively smaller effect of hypoxia on gene repression does not appear to be due to the persistence of mRNA molecules present prior hypoxia exposure. Finally, in order to get a list of core hypoxia-responsive genes we identified those that were ubiquitously expressed. To that end, we selected those genes whose expression, averaged across conditions, was detectable in at least 90% of the analyzed subsets (figure 3A labeled in purple). The resulting list included 178 genes consistently induced by hypoxia across conditions. Functional enrichment of Gene Ontology terms, indicated that core hypoxia-induced genes are mainly involved metabolic reprogramming but also in differentiation and morphogenesis, being the development of the circulatory system particularly prominent (figure 4A). On the other hand, the 113 genes consistently repressed by hypoxia across conditions, are involved in cell cycle progression, DNA replication and repair, ribosome/rRNA biogenesis and metabolism of amino acids (figure 4B). In summary, the application of a formal meta-analysis to hypoxia gene expression profiles using a random effects model lead to the identification of a set of 291 ubiquitously expressed genes whose expression is significantly altered by hypoxia by a factor of at least 0.7 log2-units. The identity of these genes can be found in supplementary table S5.

**Figure 4.**
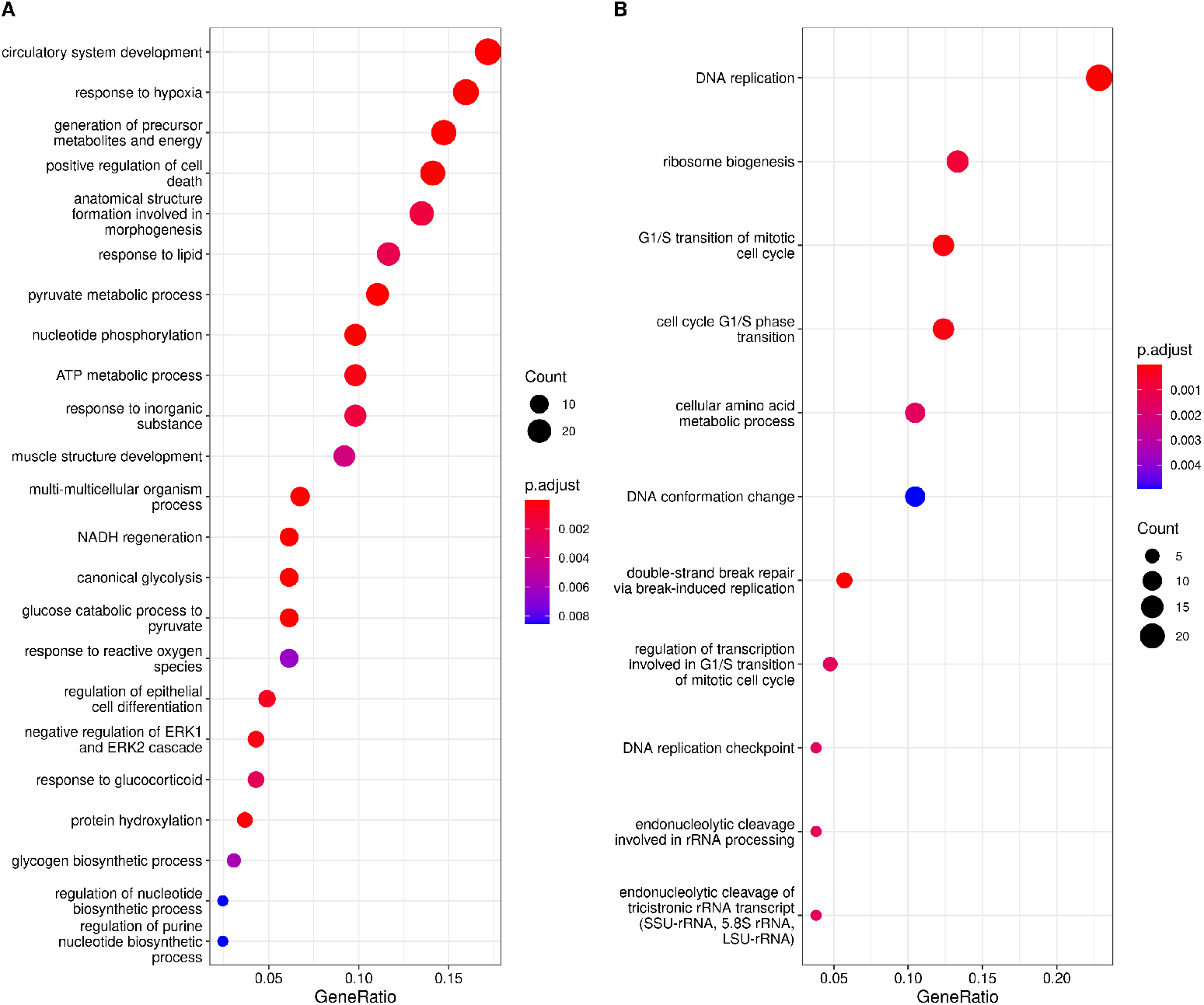
Functional enrichment analysis of the core hypoxic signature.. Gene Ontology terms significantly enriched among the core of hypoxia up- (A) and down- (B) regulated genes.

### Consistency of Meta-analysis results

To test the consistency of the pooled estimates described above, we performed a set of meta-analyses using as input all data subsets except for one and then compared the estimated effect sizes with the actual LFC observed in the subset that was left out. The process was repeated until all possibilities were exhausted. As shown in figure 5, in almost all of the cases there was a strong correlation between the pooled estimates and the actual effect sizes observed in the individual subset that was left out of the meta-analyses, with 50% of the instances showing a Pearson’s correlation coefficient over 0.82 and 75% of the cases above 0.73. Thus, the ensemble of pooled estimates it likely to predict with high accuracy the response to hypoxia in new experiments not included in the meta-analysis.

**Figure 5.**
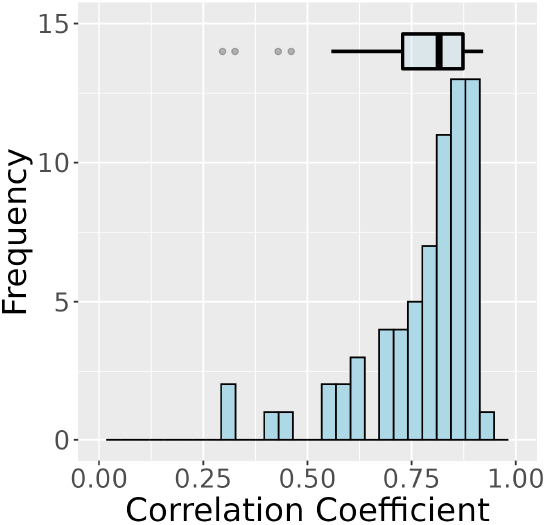
Correlation between meta-analysis’s pooled estimates and effect sizes in individual experiments.. A meta-analysis was performed for each gene using as input 63 out of the 64 subsets and the correlation between the pooled estimates and the effect sizes (Log-Fold change) in the subset that was left out of the meta-analysis was calculated. The figure represent the distribution of the Pearson correlation coefficients (Pearson’s r) observed in the 64 comparisons.

### Comparison of Meta-analyses results with a reference hypoxia signature

The core set of genes identified in the analyses described before can be considered a signature of the transcriptional response to hypoxia. Thus, we next compared the core of hypoxia-inducible genes derived from the meta-analysis with the Msig DB’s Hallmark hypoxia geneset [21], a gene signature composed of 200 genes up-regulated in response to low oxygen levels. As shown in figure 6A, the overlap between both gene sets was relatively small, with less than one third (64 out of 200) of the genes in the Hallmark hypoxia signature being present in the meta-analyses derived geneset, in spite of both genesets being similar in size, median log-fold change and nearly universal expression (supplementary table S1). Moreover, the overlap was only moderately increased when the Hallmark hypoxia signature was compared to the geneset derived from the meta-analysis without restricting to ubiquitously expressed genes (figure 6B). In order to understand the cause for the reduced overlap, we analyzed the effect of hypoxia on the expression of those genes present in the Hallmark hypoxia signature only (supplementary table S6). As shown in figure 6C, around half of the genes (50 out of 103) in this group were not included in the meta-analyses-derived signature because the pooled estimate of their induction by hypoxia was below the threshold value of 0.7, in spite of being statistically significant (labeled as “UP small ES” and shown in blue color). However, the remaining genes (53 out of 103) did not show a statistically significant induction by hypoxia (figure 6C labeled as “Non DEG” and shown in green color) or were repressed (figure 6C labeled as “DN small ES” and shown in red color). A forest plot representing the pooled estimate of the LFC for these genes shows that they cluster around the value of zero and that, in many cases, confidence interval for the point estimate is wide (Figure 6D). Altogether these results suggest that these 53 genes are either not induced by hypoxia or show a cell-type specific induction. An example of the latter is the HMOX1 whose expression is induced, repressed or left unaltered depending on the cell-type and/or experimental conditions (supplementary figure S3). Interestingly, among the genes whose regulation by hypoxia depends on the specific experimental conditions are ALDOB, GCK and LDHC that are paralogs of ALDOA, HK1 and LDHA respectively all of which are strongly and consistently induced by hypoxia.

**Figure 6.**
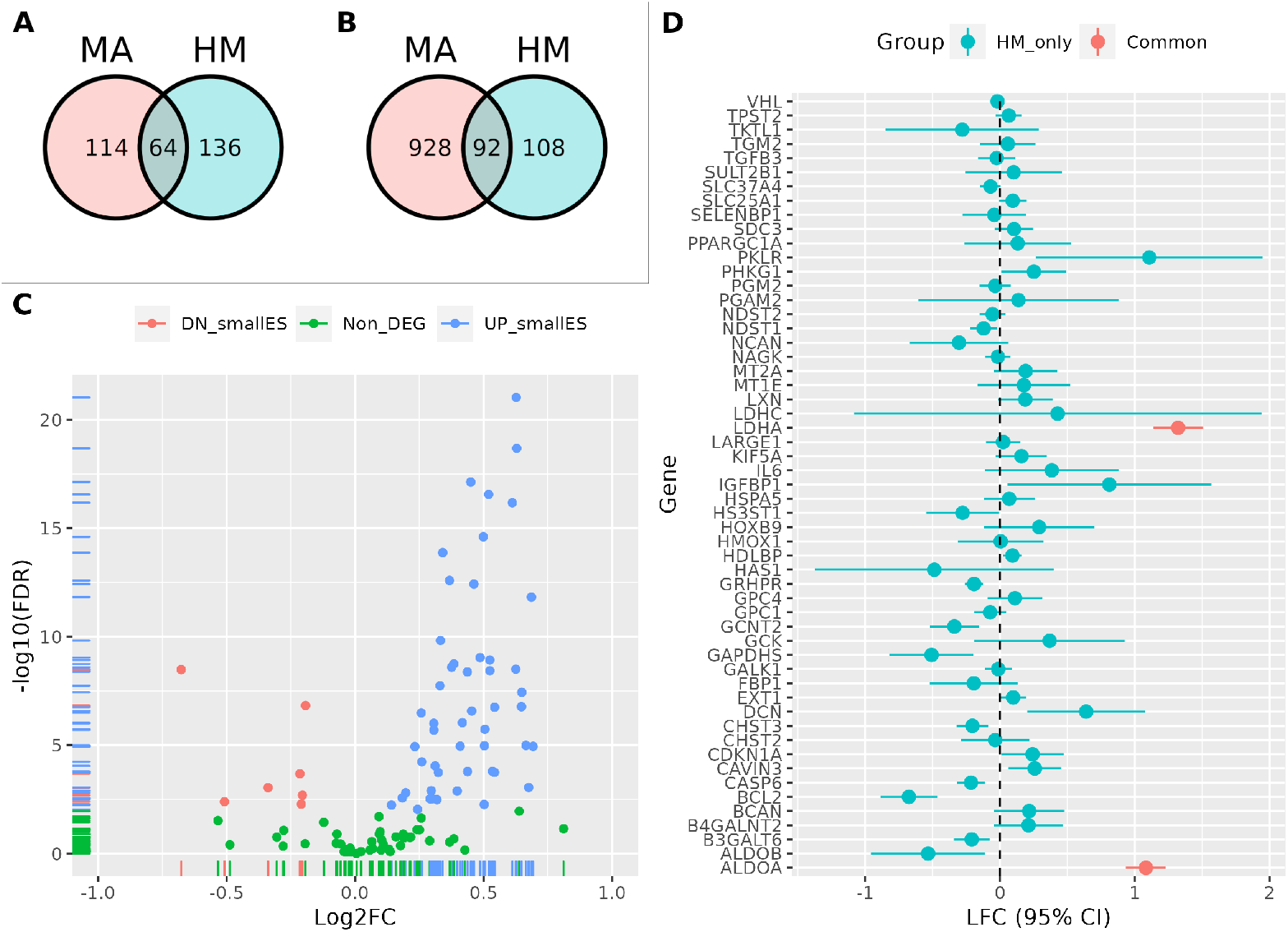
Comparison of Meta-analyses-derived and Hallmark hypoxic signatures.. Venn diagrams showing the number of genes shared by the Hallmark hypoxic signature (labeled HM) and the meta-analysis derived (labeled MA) core geneset (A), including genes expressed in over 90% of the studies, or an extended geneset including genes expressed only in some experimental conditions (B). (C) Effect of hypoxia (pooled estimates from meta-analyses) in Hallmark hypoxia signature genes not present in the meta-analyses geneset. Genes showing a statistically significant induction or repression by hypoxia are shown as blue or red dots respectively, while those whose expression is not significantly altered are shown as green dots. (D) Forest plot showing the estimated effect of hypoxia on the expression of Hallmark genes not significantly up-regulated (those labelled in red and green colors in panel C) together with the confidence interval for the point estimate. For comparison, the paralogs of ALDOB and LDHC, ALDOA and LDHA respectively, are included and labelled in red color.

Altogether, these results suggest that the meta-analyses derived gene signature improves the gene sets derived from individual studies by excluding genes whose regulation is cell type or condition specific and those with effect sizes of small magnitude.

### Cell-type specific effects of hypoxia on gene expression

The results shown above indicate that, for some genes such as HMOX1, the effect of hypoxia varies depending on the specific experimental conditions. Thus, we next studied whether cell type could influence the response to hypoxia. To that end we took advantage of the relatively large representation of endothelial cell types in the compiled dataset and performed a subgroup analysis (moderator analysis) to test the hypothesis that the differences in effect sizes were associated to the cell type used in the study. To avoid interference with other variables, we only took into consideration experiments studying gene expression in samples exposed to hypoxia for 24h and genes expressed in at least 10 non-endothelial and 7 endothelial subsets. This analysis revealed a group of 822 genes whose regulation by hypoxia was significantly different (*FDR* < 0.01) in endothelial as compared to non-endothelial cell lines (supplementary table S7), with most of the genes (712 out of 822) showing a significantly weaker regulation by hypoxia in endothelial cells.

Since many of the non-endothelial cell lines included in this analysis derive from tumors, whereas endothelial cells are primary non-transformed cells, it could be argued that the differential effect of hypoxia on endothelial is a consequence of this difference between both groups. To test this possibility, we repeated the group analysis comparing the effect of hypoxia on gene expression on three groups of samples: transformed (tumor-derived, “Tumoral”), endothelial cells (non-transformed endothelial cells, “NonT Endoth”) and other non-transformed cells different of no-endothelial lineage (non-transformed other than endothelial cells, “NonT Other”). As before, only experiments performed at 24h and genes expressed in a sufficient number of samples per group (at least 7 endothelial 10 subsets of the other two groups) were considered. This analysis identified 579 genes whose regulation by hypoxia differed significantly (*FDR* < 0.01) between groups (supplementary table ST7 Endoth Specific2). Pair-wise comparisons showed that the effect of hypoxia on gene expression was very similar for tumor cells and non-endothelial cells (Pearson’s correlation coefficient 0.903) for the 579 genes identified asdifferentially expressedbetween groups (Figure 7A). In contrast, the response to hypoxia of almost all these in endothelial cells was significantly different to that observed in the other two groups (Pearson’s correlation coefficients of 0.207 and 0.259 for the pairs NonT-Endo vs NonT-Other and NonT-Endo vs Tumoral respectively) (Figure 7A). Altogether these results indicate that there is a subset of genes whose response to hypoxia is significantly different in endothelial cells as compared to other cell types regardless their transformed status. This conclusion is supported by the heatmap in figure 7B that, by representing the effect of hypoxia on each gene (rows) across all samples (columns) sorted according to cell type, shows that the different response to hypoxia associates with the endothelial group for all clusters of genes.

**Figure 7.**
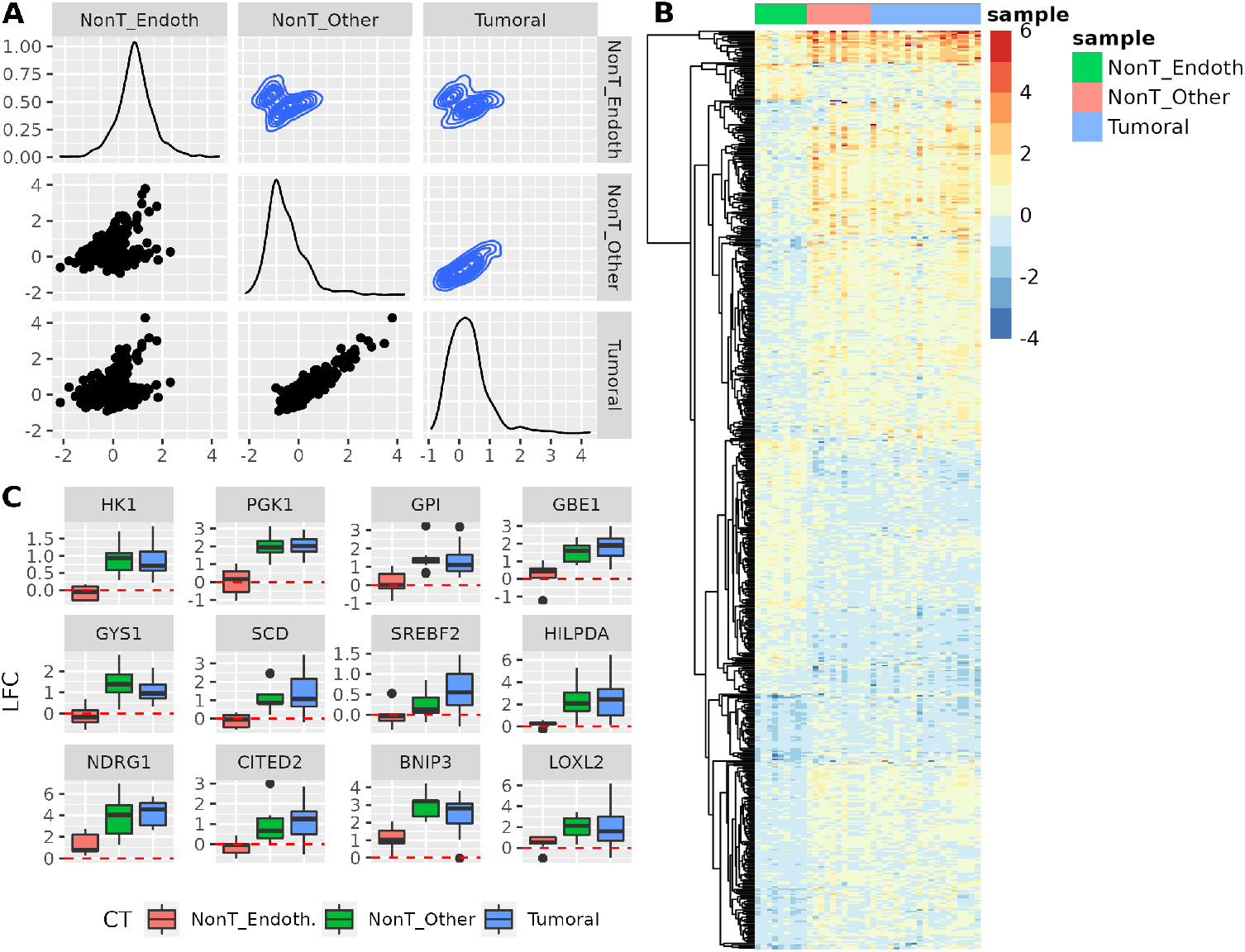
Regulation of metabolic genes is blunted in endothelial cells. A) Pairwise comparison of the effect of hypoxia across groups. The pooled effect of hypoxia was determined for each group of samples in a sub-group analysis and and their values compared to determine genes showing statistically significant differences across groups. The scatter plots show the pooled estimates (Log2-FC) for the 579 genes, each represented by a dot, showing a statistically significant response to hypoxia (*FDR* < 0.05) across groups. Each plot compares the LFC in two groups (showed on the margins of the graph). Plots in the diagonal represents the distribution of values in each sample and those in the upper half of the figure show density representation of the distribution of data points. B) Heatmap showing the effect of hypoxia on gene expression (values of LFC) in a color code as shown in the legend. The 579 differentially regulated across groups are shown in rows sorted according to their similarity in expression across samples. Columns represent samples and were sorted according to the cell type. C) The boxplots represent the distribution of the original LFC values in each group of samples for a group of selected genes. The red dotted line represents the value of zero. Differences between groups were significant in all cases (ANOVA, *FDR* < 0.05) and a post-hoc pair-wise comparison showed that the differences between endothelial cells and the other two groups were significant(Student’s t-test, *p* < 0.001), whereas the difference between the other two groups was not (*p* > 0.05).

Next we performed an enrichment analysis to identify ontology terms over-represented in the set of genes showing an altered response to hypoxia in endothelial cells. This analysis revealed that many terms related to metabolism, mostly carbohydrate catabolic pathways, were significantly (*FDR* < 0.05) over-represented as well as terms related to “response to hypoxia”. As shown in 7C, genes encoding for enzymes involved in glycolysis (e.g. HK1, PGK1 and GPI) and synthesis of glycogen (e.g. GYS1 and GBE1) as well as genes coding for proteins related to lipid metabolism (e.g. SCD, SREBF2 and HILPDA) were not induced by hypoxia in endothelial cells in sharp contrast with their consistent up-regulation in other cell types. Similarly, some typical hypoxia-regulated genes such as NDRG1, CITED2, BNIP3 and LOXL2 showed blunted induction.

Thus, group analysis identified a endothelial specific transcriptional response to hypoxia characterized by a weakened glycolytic response.

## Discussion

The integration of multiple datasets representing the transcriptional response to a given stimulus, allows for the identification of consistent changes in gene expression. However, transcriptional profiles are noisy, and correlation between different studies show poor correlation [22, 23]. Fortunately, the application of meta-analysis, appears to be a good and practical solution to reduce noise and increase signal across different studies [10]. Herein we describe the application of a formal meta-analysis procedure to identify genes whose expression is significantly modulated across a number of different gene profiling studies. This approach not only provides the identity of the genes but also a pooled estimate of the effect of the condition on the expression. Moreover, by applying a random effects model, this stategy takes into account the wide variability in gene expression expected from the integration of transcriptomes derived from different experimental conditions. The application of this approach to 69 paired normoxic/hypoxic transcriptomes representing a total of 425 samples resulted in the identification of 8556 genes, roughly 41% of the detectable genes, as significantly (*FDR* < 0.01) regulated in response to changes in oxygen tension. These results beg answering the question of the biological relevance of statistically significant but small effects of gene expression. For example, the median effect size for significantly up-regulated genes was 0.4, corresponding to a fold induction of 1.31 times over normoxia. Thus, for fifty percent of the significantly-induced genes the level of mRNA in hypoxia is at most 1.31 times higher than control levels. For some genes this small increase in expression could have important biological consequences. However, statistical significance by itself does not warrant biological relevance. For this reason we sought to identify an effect size that it is likely to have an impact of cellular biochemistry. To this end we recorded the changes in expression of genes known to have an impact on different biological processes upon exposure to hypoxia and took the median effect size value, 0.7 log2 units, as threshold. However, as only a fraction of the genes in a category are induced to initiate the biological response, this value is likely to be an underestimation. The application of the effect size threshold, together with selection of genes expressed in at least 90% of the datasets, resulted in the identification of a core set of 291 genes robustly regulated by hypoxia.

The Hallmark subset of MSigDB contains signatures generated by a computational method based on the identification of overlaps across different gene sets and retaining those genes that display coordinate expression [21]. In spite of being an invaluable and widely used resource, our results suggests that the hypoxia signature shows some shortcomings. For one thing, the signature lacks many hypoxia-regulated genes, containing only a fraction of the genes strongly and consistently regulated by hypoxia across different cell types and experimental conditions (Figure 6). The 114 core genes identified by the meta-analysis and not present in the Hallmark signature, include well characterized hypoxia-induced genes such as BCKDHA, EGLN1, several KDM family members and LOXL2 among others (supplementary table S6). On the other hand, the Hallmark signature includes some genes that are induced only in specific cell types or experimental conditions (Figure 6D) and thus, cannot be considered general hypoxia responsive genes. This result is particularly interesting as it explains the contradictory reports regarding the effect of hypoxia on specific genes such as HMOX1 [24–28] and PPARG1 [29–35] Thus, these results suggest that meta-analysis derived gene signatures are superior to other consolidated methods to identify the genes characteristic of biological processes. Given the widely use of MSigDB genesets in functional enrichment analysis, this conclusion could have far reaching implications.

Finally, the application of a formal meta-analysis approach permits the application of all associated statistical tests, including moderator analysis, to study the effect of different factors on the regulation of gene expression. Through the application of this analysis we found that endothelial cells are deficient in the induction of a relatively large set of genes in response to hypoxia. Among those genes are many enzymes involved in the metabolism of glucose, in particular those involved in glycolysis and synthesis of glycogen. It is known that HIF1A, but not EPAS1, is responsible for the hypoxic induction of glycolytic genes [36], it is tempting to speculate that the specific pattern of expression observed in endothelial cells could be a consequence of the relative importance of EPAS1 over HIF1A isoform in this cell type. In contrast to the case of endothelial cells, moderator analysis did not identified large differences in the response to hypoxia comparing normal and transformed cells. This result suggest that, in general, oncogenic transformation does not substantially affect the transcriptional response to hypoxia.

Altogether our results suggest approaches based on formal meta-analyses are powerful method to identify universal as well as condition-specific gene expression patterns.

## Supporting Information

## Acknowledgments

This work was supported by Grants SAF2017-88771-R and PID2020-118821RB-I00 funded by MCIN/AEI/10.13039/501100011033 and by “ERDF A way of making Europe” and by grant IND2019/BMD-17134 funded by Autonomous Autonomous Community of Madrid.

## Supporting Information Legends

**Table S1.** Comparison of Meta-analysis derived core genes and Hall Mark hypoxia signature. MA, Meta-analysis; HM, MSig’s Hypoxia Hall Mark; Size, number of genes in the signature; Ub., Number of genes present in more than 90% of the analyzed subsets; 1QLFC, Log2 Fold Change first quartile;1QLFC, Log2 Fold Change third quartile.

| Signature | Size | Ub. | 1QLFC | 3QLFC |
| --- | --- | --- | --- | --- |
| MA | 178 | 178 | 0.76 | 1.33 |
| HM | 200 | 195 | 0.67 | 1.14 |

**Table S2. *Studies selected from database search.*** *GSE_ID, GEO Series ID; The oxygen tensions hypoxia exposure times (in hours) and cell type(s) included in the study are shown in columns O2, Hx_time and CT respectively.*

**Table S3. *Metadata of studies kept after filtering original datasets.*** *Columns A-P correspond to the information associated to samples in the NCBI’s Sequence Read Archive (SRA).GSE_ID, GEO Series ID; The oxygen tension hypoxia exposure time(in hours) and cell type and transformed status of each sample is recorded in columns O2, Hx_time, Cell type/CT and transformed respectively.*

**Figure S1.**
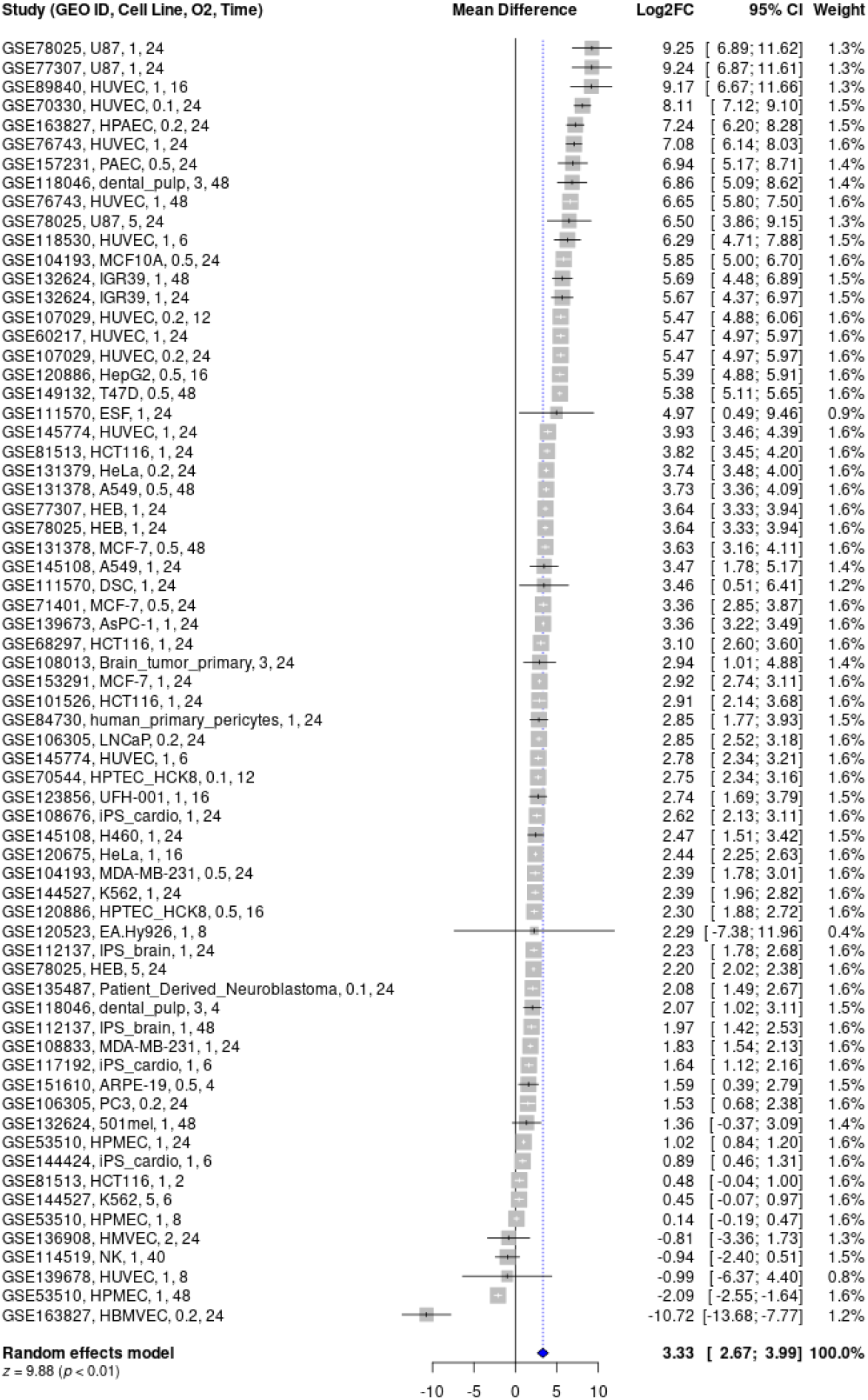
Meta-analysis of the effect of hypoxia on EGLN3 gene expression. Forest plot representing the individual effect size (LFC) estimates as grey boxes whose size is inversely proportional to the precision of the estimate. Each estimate derives from the study indicated on the left column ()“Study”), which includes information about GEO record and experimental conditions: cell line, concentration of oxygen (percentage of oxygen in gas mixture) during exposure to hypoxia and length of hypoxia in hours. The pooled effect size estimate is shown at the bottom as a blue diamond. The columns on the left indicate the values of the effect size (Log2FC), the 95% confidence interval for the estimate and the weight of each observation on the pooled effect.

**Figure S2.**
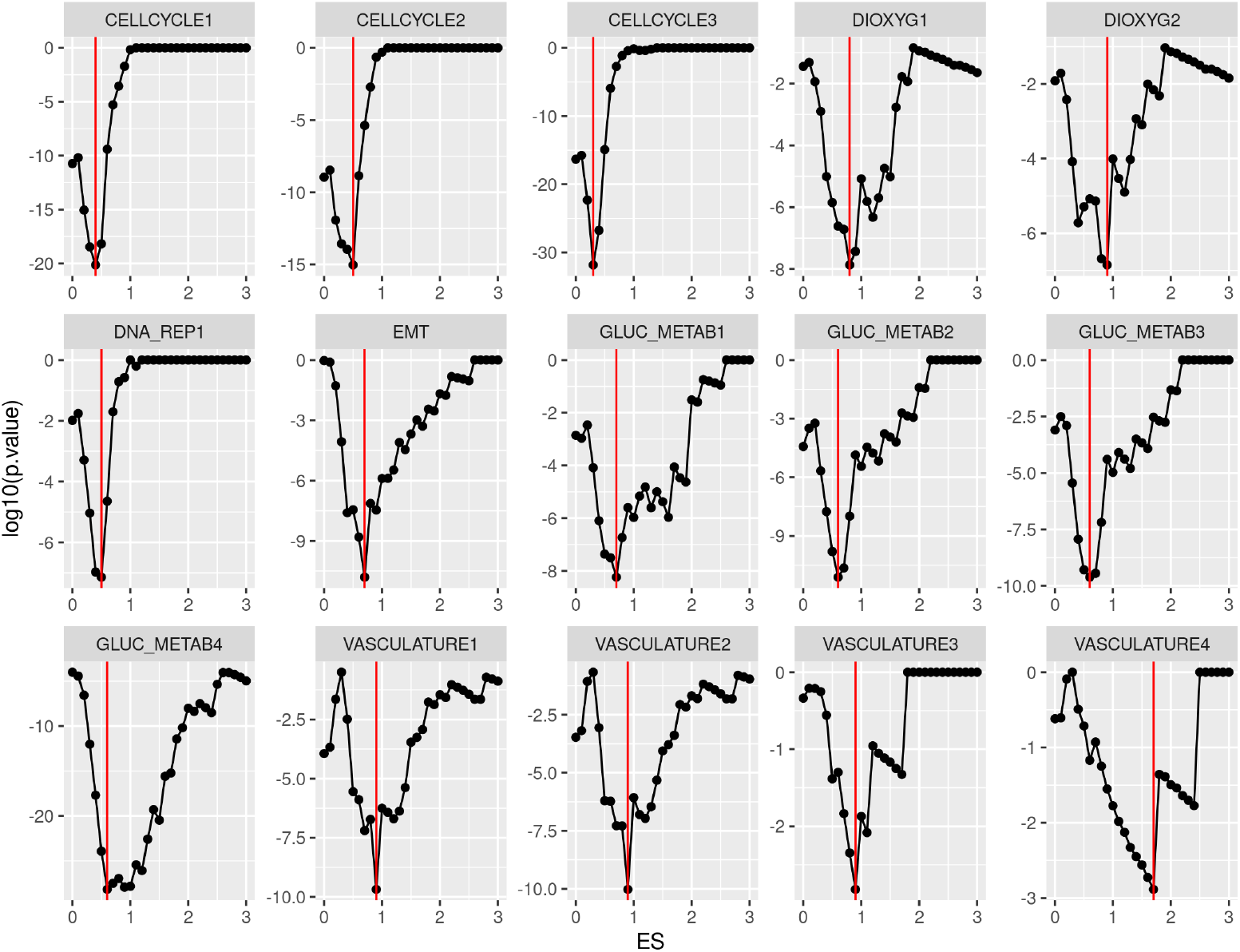
Identification of an effect size value that maximizes the association between gene expression and cellular responses to hypoxia.. Genes showing an statistically significant change in expression (*FDR* < 0.01) of a magnitude (*Log*2*FC*) above the indicated values (x axis) were categorized as differentially expressed in response to hypoxia. Then, genes were further classified into mutually exclusive subgroups according to their membership regarding the indicated biological processes. Finally, the association between the response to hypoxia and the biological process was assessed using a Fisher’s exact test. Each graph shows the p-value for the the association between the indicated biological process (Gene Ontology Biological Processes: “DNA DEPENDENT DNA REPLICATION” (CELLCYCLE1), “CELL CYCLE DNA REPLICATION” (CELLCYCLE2),; MSig DB Hallmark: “G2M CHECKPOINT” (CELLCYCLE3),…) and the response to hypoxia at different Log2 FC cut-off values.

**Table S4. *Compiled Meta-analyses results.*** *Gene_symbol, Human Genome Nomenclature gene symbol; N, number of Nx-Hx sample pairs were gene expression was detectable; G.random, pooled estimate of effect size (log2 Hx/Nx); seG.random, Standard Error of the pooled estimate; lciG.random and uciG.random, lower and upper 95% confidence intervals for the pooled estimate; pval.random; estatistical signicance for the pooled estimate being equal to zero; ajdP.random, FDR-adjusted p-values.*

**Table S5. *Effect of hypoxia on the expression of the genes identified in the meta-analysis as the core hypoxic signature.*** *Columns A-E as in table S4. Remaining columns indicate the effect of hypoxia (log2 Hx/Nx) on the gene indicated in column Gene_Symbol as determined in the study indicated in the column name. Study column names contain the following four pieces of information: GSE ID, cell type, exposure time (in hours) and percentaje of oxygen sepparated by underscores.*

**Figure S3.**
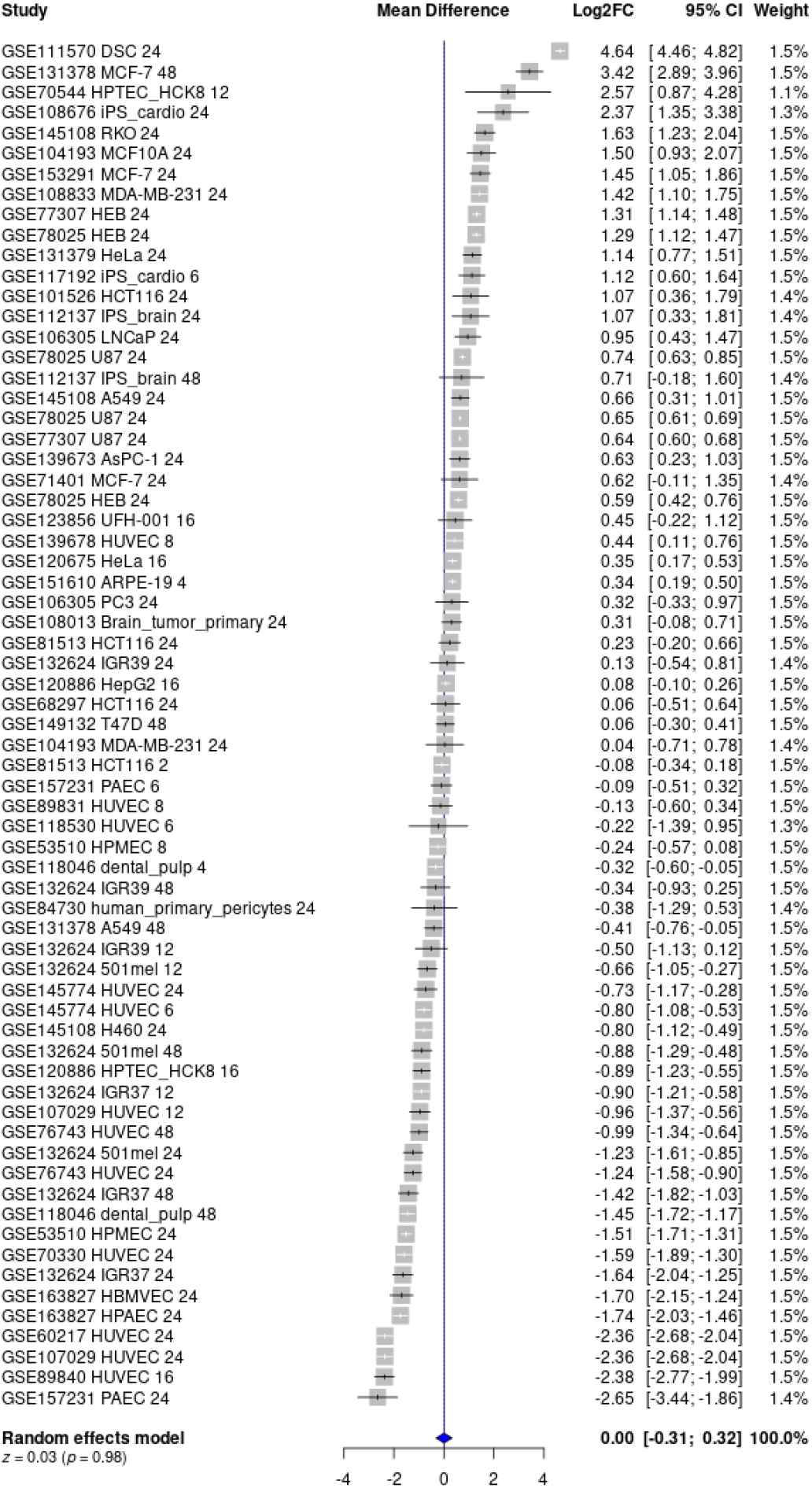
Meta-analysis of the effect of hypoxia on HMOX1 gene expression. Forest plot representing the individual effect size (LFC) estimates as grey boxes whose size is inversely proportional to the precision of the estimate. Each estimate derives from the study indicated on the left column (“Study”), which includes information about GEO record and experimental conditions: cell line, concentration of oxygen (percentage of oxygen in gas mixture) during exposure to hypoxia and length of hypoxia in hours. The pooled effect size estimate is shown at the bottom as a blue diamond. The columns on the left indicate the values of the effect size (Log2FC), the 95% confidence interval for the estimate and the weight of each observation on the pooled effect.

**Table S6. *Effect of hypoxia on the expression of the genes included in gene signatures.*** *Columns A,D-G as in table S4. Columns MA and HM indicate whether the gene (row) is included in the Meta-analysis and MSigDB’s Hall Mark gene signatures respectively.*

**Table S7. *Genes whose regulation by hypoxia is significantly different in endothelial cells compared with the rest of cell types.*** *Gene_symbol, Human Genome Nomenclature gene symbol; N_yes and N_no, number of Endothelial and not-endothelial Nx-Hx sample pairs respectively were gene expression was detectable; MD_yes and MD_no, pooled estimate of effect size (log2 Hx/Nx) for the endothelial and non-endothelial groups of samples; pval.random; estatistical signicance for the difference between groups being equal to zero; ajdP.random, FDR-adjusted p-values.*

**Table S8. *Genes whose regulation by hypoxia is significantly different in endothelial cells compared with the other non-transformed and transformed cell types.*** *Columns as in table S4. Suffixes “e”, “o” and “t” refer to endothelial, other-non transformed and transformed cells respectively.*

## Notes

### Competing Interest Statement

The authors have declared no competing interest.

